# Cathartocytosis: Jettisoning of Unwanted Material during Cellular Reprogramming

**DOI:** 10.1101/2024.06.11.598489

**Authors:** Jeffrey W. Brown, Xiaobo Lin, Gabriel Anthony Nicolazzi, Xuemei Liu, Thanh Nguyen, Megan D. Radyk, Joseph Burclaff, Jason C. Mills

## Abstract

Injury can cause differentiated cells to undergo massive reprogramming to become proliferative to repair tissue via a cellular program called paligenosis. Gastric digestive-enzyme-secreting chief cells use paligenosis to reprogram into progenitor-like Spasmolytic-Polypeptide Expressing Metaplasia (SPEM) cells. Stage 1 of paligenosis is the downscaling of mature cell architecture via a process involving lysosomes. Here, we noticed that sulfated glycoproteins were not only digested during paligenosis but also excreted into the gland lumen. Various genetic and pharmacological approaches showed that endoplasmic reticulum membranes and secretory granule cargo were also excreted and that the process proceeded in parallel with, but was mechanistically independent of autophagy. 3-dimensional light and electron-microscopy demonstrated that excretion occurred via unique, complex, multi-chambered invaginations of the apical plasma membrane. As this lysosome-independent cell cleansing process does not seem to have been priorly described, we termed it “cathartocytosis”. Cathartocytosis allows a cell to rapidly eject excess material without waiting for autophagic and lysosomal digestion. We speculate the ejection of sulfated glycoproteins would aid in downscaling and might also help bind and flush pathogens away from tissue.

## INTRODUCTION

When injury causes loss of mature cells in tissues with constitutively active, multipotent stem cells, the tissue can be repaired by increasing stem cell proliferation and differentiation thereby replacing both the lost cell mass and maintaining the stem cell pool (Meyer et al., 2022). However, when stem cells themselves are damaged or when tissues lack dedicated stem cells, the burden of repair falls on mature, differentiated cells (Brown et al., 2022). As a consequence, differentiated cells have evolved a capacity to respond to injury by reprogramming to a downscaled proliferative phenotype to regenerate damaged and dead cells and maintain barrier function (Cho et al., 2024). It has been proposed an evolutionarily conserved program executes reprogramming of mature cells to cycling progenitors. Specifically, our group has shown that following injury, post-mitotic differentiated cells can reprogram into proliferative, immature-appearing cells through a series of three, stepwise stages in a conserved process known as paligenosis (Willet et al., 2018).

Briefly, during the response to injury, a cell undergoing paligenosis tunes down mTORC1 activity and massively upregulates autodegradative machinery characterized by expanded lysosome activity and induction of autophagy (Stage 1; (Radyk et al., 2021)). During this phase of cellular downscaling, mature cellular machinery is dismantled and removed, becoming superfluous as the cell transitions to a proliferative progenitor state for tissue repair. In Stage 2 of paligenosis, autophagy subsides, and mTORC1 activity increases coincident with the expression of metaplasia-related, progenitor-cell proteins like SOX9 (Willet et al., 2023). Next, cells can proceed into the cell cycle (Stage 3) if they successfully suppress p53 and its licensing function (Miao et al., 2021; Miao et al., 2024). The reprogrammed chief cells are known as Spasmolytic Polypeptide Expressing Metaplasia (SPEM) cells (Nomura et al., 2004; Schmidt et al., 1999; Wang et al., 1998; Yamaguchi et al., 2002a, b) and the organ level metaplastic transformation referred to as pyloric or pseudopyloric metaplasia due to the antralized appearance of the corpus with light microscopy (Goldenring, 2018; Goldenring and Mills, 2022).

Here, we further examine how mature, differentiated cells downscale their cellular machinery after injury and uncover a previously uncharacterized cellular process to extrude large amounts of cell material through the apical membrane that occurs coincident with, but mechanistically independent from, autophagy. We structurally and functionally characterize this process and term it cathartocytosis [Greek: cellular cleansing].

We show cathartocytosis is associated with the expression and secretion of acidic (sialylated & sulfated) glycoproteins, which have long been recognized as a histologic feature of gastric metaplasia (a pre-cancerous lesion) and cancer in the esophagus, stomach, and pancreas (Das and Brown, 2023; Das et al., 2021; Hakkinen et al., 1968; Hakkinen and Viikari, 1969; Piazuelo et al., 2004). More recently, we have shown that extracellular (presumably secreted) sulfated glycoproteins can be used as a clinical biomarker to detect these cellular transformations (Das et al., 2019; Das et al., 2014). Such transitions, and associated carcinogenesis, have been documented in the diagnostic pathology literature on gastric cancer where the absence or presence of particular acidic mucins in columnar cells defines different subtypes of metaplasia and can help stratify risk for gastric adenocarcinoma development (Das and Brown, 2023; Shah et al., 2020).

Here, we carefully track the subcellular distribution and movement of both the sulfated mucins in the zymogenic granules and the endoplasmic reticulum as the gastric chief cells downscale their cellular machinery *en route* to metaplasia. We find that these two organellar compartments, which comprise the vast majority of the gastric chief cell cytoplasm, are not only routed to autophagic degradation in Rab7^+^/Lamp2^+^ late endosomes / lysosomes (LE/Ly), a trafficking mechanism that has been previously described, but also excreted. We find that this excretion is mechanistically independent of canonical autophagy. Using focused ion beam scanning electron microscopy (FIB-SEM), we structurally characterize the secretory apparatus for the excretion process and find it occurs via phagophore-shaped apical membrane invaginations. We propose the term cathartocytosis [Greek: Cellular Cleansing] to describe this excretion process and believe it is at least partially responsible for delivery of sulfated mucins and extracellular vesicles into the lumen of the gastric gland. As these sulfated mucins are specific biomarkers in patients to assay for metaplasia and cancer, cathartocytosis may have clinical implications.

## RESULTS

### Sulfated glycoproteins are expressed in gastric epithelium in homeostasis and after metaplastic injury

As mucins are known to stage precancerous lesions and cancer in gastrointestinal epithelia, we wanted to better characterize the expression, distribution, and secretion of sulfated glycoproteins during metaplastic transition in the stomach. We used the antibody Das-1, which recognizes sulfated glycosylation epitopes as they arise during oncogenic transformation in the foregut (i.e., stomach, esophagus, and pancreas)(Das and Brown, 2023). Indeed, Das-1 immunolabeling is a sensitive and specific biomarker for foregut metaplasia, dysplasia, and cancer (Bodger et al., 2003; Das et al., 2021; Das et al., 2019; Das et al., 2014; Das et al., 1994; Piazuelo et al., 2004). Das-1 specifically recognizes 3’-Sulfo-Le^A^ and 3’-Sulfo-Le^C^ (Brown et al., 2021) as epitopes (Supplemental Figure 1). To determine if the sulfated glycotopes recognized by Das-1 were present on large molecular weight proteins or lipids, we analyzed Das-1 bands on western blot from a human gastrointestinal cell line LS174T, which are a colorectal cell line but with a mutational profile that more resembles that of transformed foregut epithelial cells (K-Ras^G12D^; PIK3CA^H1047R^; E-cadherin-negative) (Ahmed et al., 2013; Efstathiou et al., 1999). The epitope was largely removed by treatment with NaOH but was unaffected by PNGase F (removes N-linked glycans), O-Glycosidase (removes linear, unbranched O-linked disaccharides from glycoproteins), or Neuraminidase (which cleaves linear and branched terminal sialic acids) (Supplemental Figure 2). Because O-Glycosidase can remove only unbranched O-linkages, while NaOH removes all O-linked glycans, these data suggest that the Das-1 epitope is largely present on branched, O-linked glycan trees. Note that PNGaseF caused a shift in mobility but not in Das-1 labeling, suggesting the presence of some N-Linked glycans elsewhere on the proteins that carry the Das-1 epitopes such that the loss of the N-linked moieties causes the band to run faster but does not affect Das-1 binding. In other words, because the immunoreactivity remained, those glycans must be on separate glycan trees without terminal 3’-Sulfo-Le^A/C^ groups such that they contribute only to the total mass of the glycoprotein. We have also excluded the possibility of the Das-1 epitope being present on glycolipids as it was in the pellet and not the supernatant following chloroform extraction (Supplemental 2D-E) and reactivity remains even after xylene processing of murine tissue (Supplemental Figure 2F; note both chloroform and xylene extract lipids). Altogether, the results show that Das-1 epitope recognizes 3’-Sulfo-Le^A^ and 3’-Sulfo-Le^C^ attached via branched O-linkage to complex glycoproteins.

Mice are unable to express 3’-Sulfo-Le^A^ because that structure depends on α(1-4) fucosylation, and the requisite fucosyl transferase *Fut3* is a pseudogene in mice (Gersten et al., 1995); thus, 3’-Sulfo-Le^C^ is the Das-1 murine antigen (Supplemental Figure 1). NaOH treatment on mouse stomach lysates also removed the Das-1 epitope (Supplemental Figure 2F-G). Thus, the results in murine stomach tissue and with glycoprotein stripping experiments in blots from human cells are consistent with Das-1 recognizing glycoproteins bearing branched O-linked glycan trees terminating in 3’-Sulfo-Le^A^ and 3’-Sulfo-Le^C^ moieties. Das-1 immunohistochemistry (IHC) showed that, unlike the normal *human* stomach (Brown et al., 2021) where Das-1 reactivity is absent at homeostasis, homeostatic murine chief cells express sulfated glycoproteins (Figure 1A; Supplemental Figure 3). The subcellular distribution was vesicular and showed consistent proximity to and/or colocalization with, the GIF+ secretory granules in cell apex (i.e. luminal to the nucleus (Supplemental Figure 3)).

**Figure 1.**
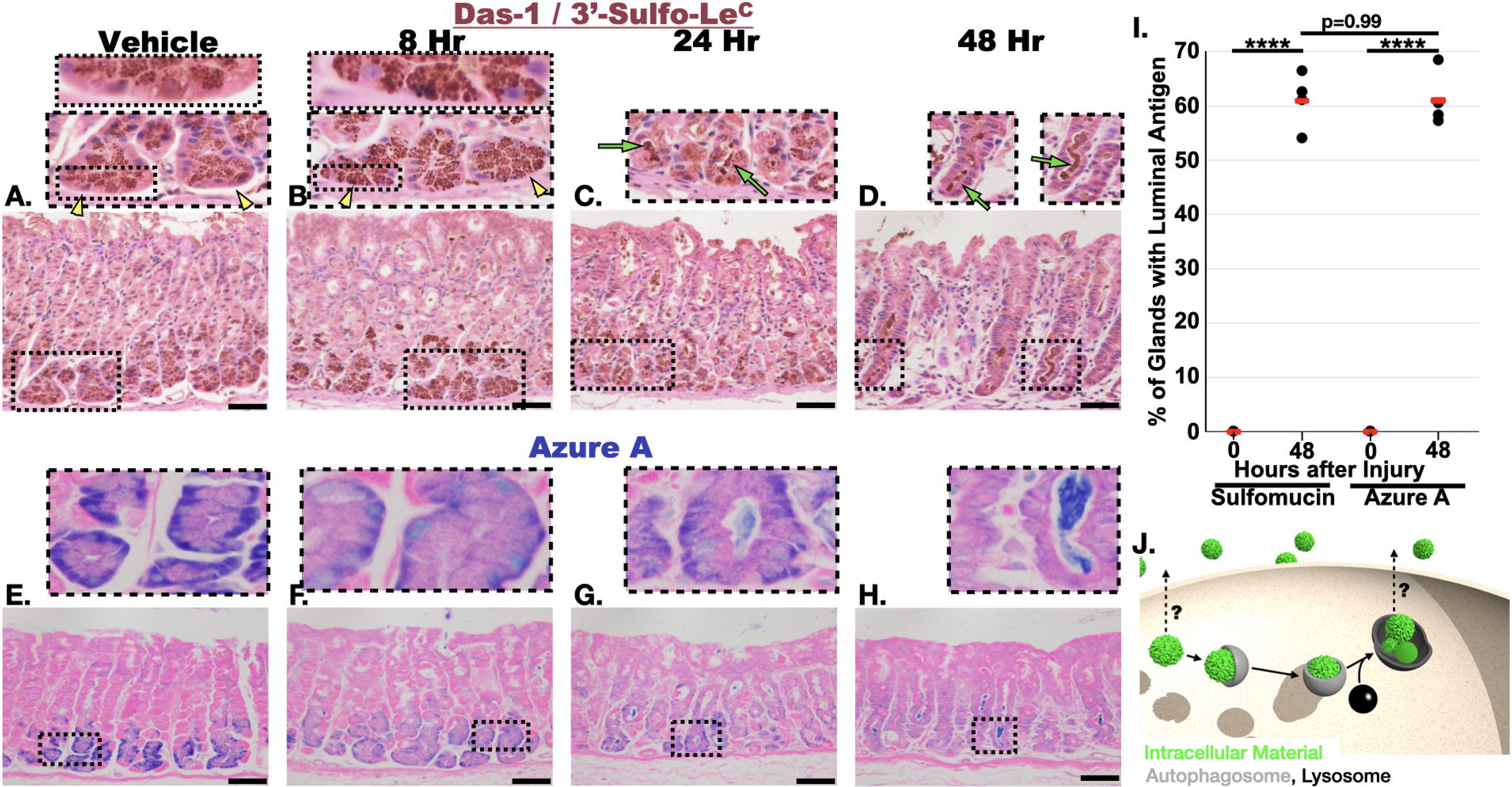
Cellular material is excreted during paligenosis. **A-D.** Micrographs of the mouse stomach after treatment with vehicle **(A, E)**, and after injury (8 hrs **(B, F)**, 24 hrs **(C, G)**, and 72 hours **(D, H)** following the first injection of high-dose tamoxifen). **A-D.** are immunostained with Das-1, which recognizes the glycosylation epitope 3’-Sulfo-Le^A/C^ (Brown et al., 2021), **E-H** stained with Azure A and Eosin**. I.** Quantitation of secreted luminal antigens 48 hours after injury. Mean of all mice (n=4-5) per condition depending on experiment. Pairwise significance calculated with T-Test. **J.** Schematic depicting potential trajectories for secreted of generic cytoplasmic organelles (green: Intracellular material like granule, ER, mitochondria, etc.). Grey: classical autophagy; Black: Lysosome. Yellow Arrowheads demonstrate the loss of apical compartmentalization of the zymogenic granules 8 hours after injury, relative to vehicle. The secretory granules are apical to the nucleus in A, but also basal to the nucleus in B. Green arrows identify secreted sulfated glycoproteins. Scale Bars = 50 µm.

We next sought to determine the effects of injury-induced reprogramming on sulfated glycoproteins. Injection of high doses of tamoxifen causes a characteristic, reversible injury pattern in the gastric epithelium marked by loss of acid-pumping parietal cells and reprogramming of chief cells into a metaplastic lineage. The effects of tamoxifen are independent of estrogen modulation and occur in both sexes (Huh et al., 2012; Keeley et al., 2019; Saenz and Mills, 2018). The conversion of the large secretory architecture of a chief cell into downscaled, mitotic SPEM cell occurs by a stereotypical sequence of molecular-cellular events known as paligenosis (Brown et al., 2022; Radyk et al., 2021; Willet et al., 2018) and can be completed in as little as 48 hours from first injection of tamoxifen.

The first stage of paligenosis lasts up to 24 hours and involves massive upregulation of autophagic and lysosomal structures as cells remodel their secretory apparatus. We noted some relocalization basally of Das-1-labeled vesicles by 8 hours (Fig. 1B); however, by 24 hours, a timepoint when injured chief cells are completing the lysosomal/autophagic degradation stage and the cell has shrunk considerably, we observed dramatic reorganization of Das-1 labeled structures. Namely, sulfomucins began to appear in the lumen of the gland apical to chief cells (Figure 1C). By 48 hours, a time point when the majority of cells have converted to the proliferative SPEM phenotype, the sulfated glycoproteins were no longer detected in the downscaled chief cells. Instead, they could be found as cast-like structures throughout the lumens of gastric glands (Figure 1D).

### Extrusion of cellular material and organelles during downscaling in paligenosis

The elaborate architecture of a differentiated gastric chief cell is characterized by a basal, globoid nucleus surrounded by profusely abundant, tightly-packed lamellar rough ER (rER) with an apical collection of large secretory granules (Figure 1A, 1E, 2A, Figures S3-5). We have previously studied how lysosomes and autophagosomes are induced during paligenosis Stage 1 (Radyk et al., 2021). Here, we note massive rearrangement and decrease of the rER during these same first 24 hours. We followed rER downscaling using focused ion beam scanning electron microscopy (FIB-SEM), the histological stain Azure A (Figure 1E-H, Figure S4), and two different antibodies against the ER resident proteins (1) peptide protein disulfide isomerase (PDI) and (2) Translocon-associated protein alpha subunit (TRAP alpha) (Figure S5). All techniques yielded consistent patterns: dense staining in the base of the cell, becoming more muted apically where zymogenic granules fill most of the cytoplasm, appearing as punched out holes in a more lattice-like ER pattern (Figure 1, Figure 2A, Figure S4, Figure S5). Azure A allowed easy ER visualization using a simple histological stain and light microscopy (Figure 1E-H, Figure S4), while the ER is not as distinct with anti-PDI on IHC. Thus, visualizing PDI or TRAP-alpha generally requires confocal microscopy with optical sectioning to obtain high-resolution images (Figure S5).

**Figure 2.**
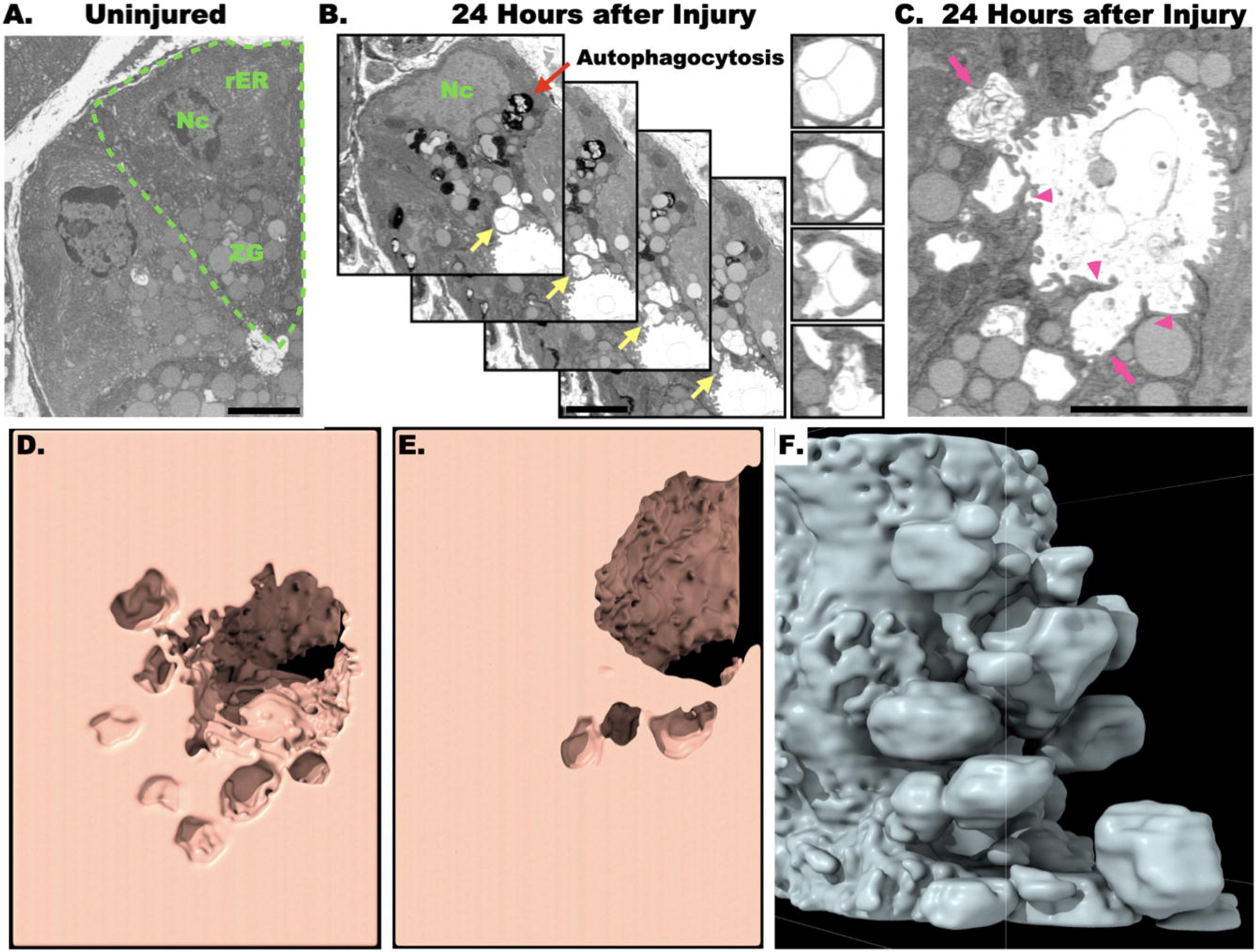
Focused ion beam scanning electron microscopy (FIB-SEM) demonstrates apical membrane deformation and secreted vesicular material 24 hours after injury. **A.** Vehicle-treated mouse stomach displaying two, mature gastric chief cells in two-dimensional electron microscopic section. The boundaries and organellar compartments of ones of these cells is annotated in green (Nc is nucleus; rER is rough endoplasmic reticulum; ZG are zymogenic granules). **B.** Four sequential FIB-SEM micrographs depicting excretion of large, electron-lucent extracellular vesicles and membranous material (red arrow: lysosomes and various stages of autophagosomes; yellow arrows highlights apical plasma membrane shown in inset at right). **C**. Another section from FIB-SEM showing membrane material being extruded apically (upper arrow) as well as the gland lumen. Note distorted projections of apical plasma membrane into lumen (magenta arrowheads) and pit-like invaginations of apical plasma membrane (magenta arrowheads). **D-F.** Three-dimensional reconstructions produced by stacking FIB-SEM sections. **D, E.** Two slices through a three-dimensional reconstruction of how the gland lumen invaginates into an interconnected network of saccules and channels made by the plasma membrane of a paligenotic chief cell. The entire clipping series is available in Supplemental Movie 1. **F.** Similar view but with convex reconstruction instead of concave. Note that the paligenotic chief cell demonstrates a series of communicating chambers invaginating into apical cytoplasm. A 360° rotational series of this model is provided as supplemental movie 2. Scale Bars = 4 µm.

Azure A is most commonly used in thin liquid chromatography to identify sulfatides (sulfated glycolipids) (Kean, 1968) as well as sulfated glycosylaminoglycans (Berman et al., 1971). Though we do not know the exact macromolecules bound by Azure A in gastric epithelium, we hypothesize they are sulfated glycosylaminoglycans (GAGs) rather than sulfated glycolipids because (1) the tissue was processed with xylene, which removes lipids and (2) antigen retrieval results in loss of Azure A material (unlike proteins, GAGs are not crosslinked with formalin (Kiernan, 2000)). We also observed staining of nuclei by Azure A, likely due to association with acidic deoxyribonucleic (Figure S6); however, nuclei had a distinct lighter, royal blue staining compared to the presumptive ER (Figure 1E-H, Figure S4).

Following injury, the dense basal staining of homeostatic cells was first consistently and notably decreased by 8 hours (Figure 1F, Figure S4B,F). Analogous to the sulfated mucins we tracked using Das-1, extrusion or secretion of Azure A-stained material occurred at 24 hours, and the gland lumens were full of ER markers by 72 hours (Figure 1G,H, Figure S4, Figure S5). By 48 hours, ∼60% of gastric glands had luminal Azure A and Das-1 antigen (Figure 1I). As we observed paligenotic cells extruding or secreting an organelle (namely, endoplasmic reticulum) along with sulfated glycoproteins, the phenomenon was clearly different from normal secretion in chief cells where cargo (namely digestive enzymes) is elaborated extracellularly by apocrine secretion.

### Changes in the apical plasma membrane during excretion

To gain ultrastructural insight into cellular changes during paligenosis, we used FIB-SEM to study the three-dimensional architecture of chief cells at baseline and after induction of metaplastic downscaling with tamoxifen. We chose to analyze the 24-hour time point because at this point the cells are (1) completing their massive downscaling (Radyk et al., 2021; Willet et al., 2018) and (2) because that is when we observed both residual intracellular and newly arising extracellular glycoproteins and endoplasmic reticulum contents (Figure 1C, G; Figure S4C, G). Consequently, active organellar extrusion is occurring at the 24-hour time point. As expected, the FIB-SEM serial sections showed that a large portion of the downscaling cytoplasm harbored various stages of autophagosomes and lysosomal structures along with rER that had already shrunk considerably from homeostasis (Figure 2A, B). Residual secretory granules were also present (Figure 2B). Note that during downscaling, the cells adopt a more cuboidal cellular shape with retraction of the apical membrane towards the cell base. As apical membrane retracts basally, the glandular lumen opens, correlating with expansion of the luminal apical plasma membrane (Figure 3C,E). These changes are hallmarks of paligenosis of chief cells (Radyk et al., 2021); however, we also noted previously uncharacterized, extensive irregularities in the apical cell membranes, including invaginations harboring vesicular-membranous material of different sizes (Figure 2C). When the membranous material appeared in a vesicle-like morphology, the lumens of such structures were electron-lucent, similar to the extracellular lumen of the gland. Both the electron lucency and highly variable sizes of these excreted membranous structures would be atypical for exosomes released by fusion of multivesicular bodies with the apical membrane.

**Figure 3.**
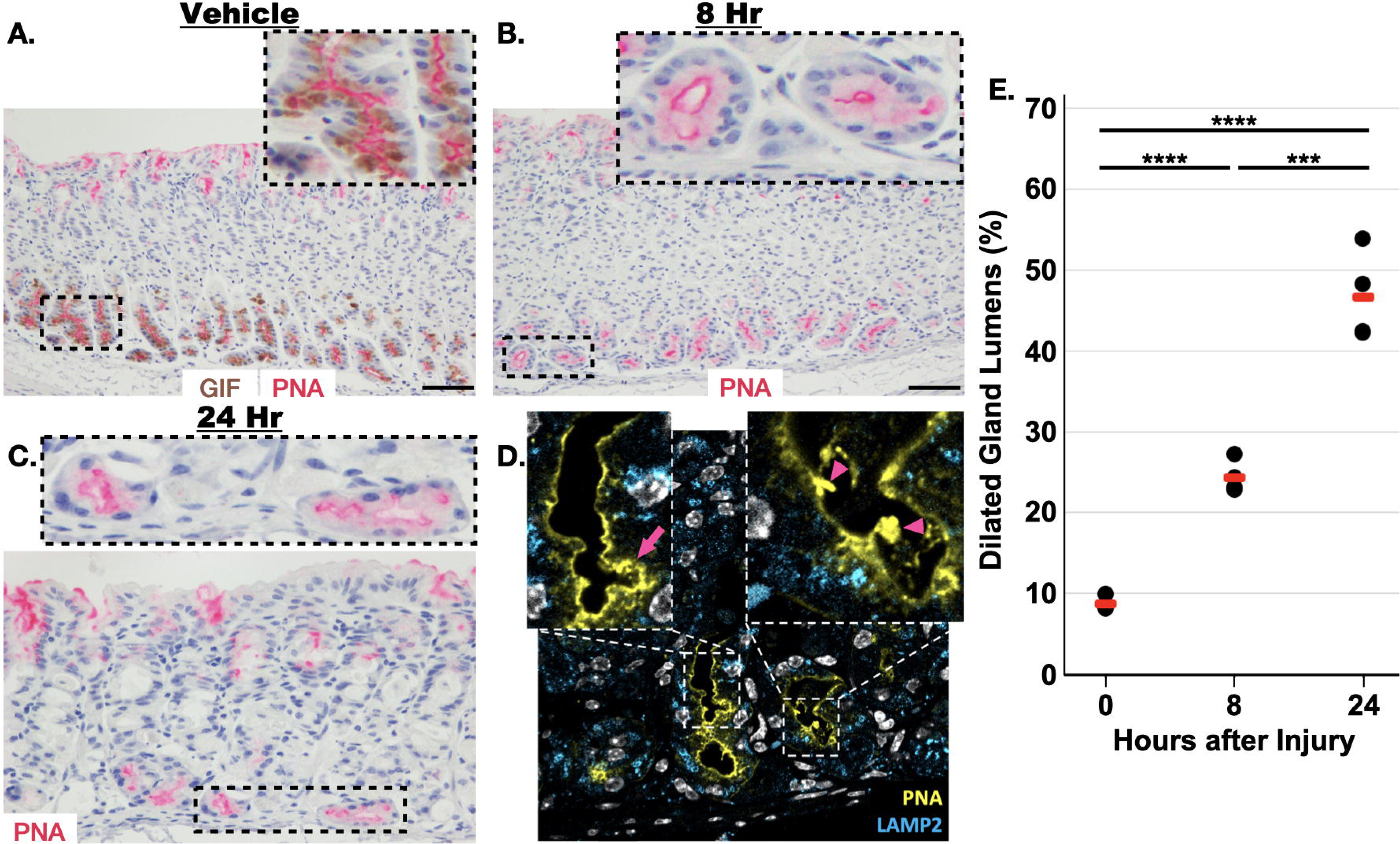
Immunohistochemical and confocal immunofluorescence highlights apical membrane distortion in stage 1 of paligenosis. **A.** PNA Lectin (magenta) is specifically reactive to the apical membrane of chief cells, which can be identified by anti-GIF antibody (brown). **B**. 8 hours after injury, the gland lumens begin to dilate. **C.** At 24 hours, the apical membrane of the gastric chief cell becomes convoluted. **D**. Using optical sectioning, these convolutions are due to invaginations (magenta arrows) and flaps of membrane (magenta arrowheads), as seen ultrastructurally in Figure 2C. Pseudocoloring: White: Nucleus (DAPI); Yellow: PNA Lectin; Blue: Lamp2.Scale bars in A-B = 100 µm. Scale bar in C = 50 µm. **E.** Quantitation of gland dilation as a function of time after injury. Black Dots= mean fraction per individual mouse from > 200 fundic glands per mouse, Red Line: Mean of all mice (n=4) per condition depending on experiment. Significance calculated with one-way ANOVA of means with Tukey’s multiple comparison test.

Segmentation (i.e., 3-dimensional reconstruction) of the cell beginning at the slice depicted in Figure 2C showed that the electron-lucent regions in the apical cell cytoplasm were actually not enclosed within the cytoplasm but rather topologically part of a network of chambers all emptying into the gland lumen (Figure 2D-F, Supplemental Movie 1). The process was apical-specific with no detectable irregularities noted in the basolateral plasma membrane. We also segmented and rendered a view of the gland lumen from a convexity perspective, where again it can be appreciated how the apical plasma membrane dives inwardly into multichambered, interconnected cellular compartments (Figure 2F, Supplemental Movie 2).

We next studied the apical invaginations at light-microscopic resolution using the lectin Peanut Agglutinin (PNA) that specifically highlights the chief cell apical plasma membrane (Figure 3A) (Falk et al., 1994; Gomez-Santos et al., 2021; Okamoto and Forte, 1988). The PNA staining highlighted the downscaling of chief cells and widening of the gland lumen beginning at 8 hours after high-dose tamoxifen (Figure 3B). At 24-hours, cell downscaling was present throughout the gland base and we observed convoluted apical plasma membrane similar to that present in FIB-SEM (Figure 3C). Optical sectioning using confocal microscopy further highlighted the apical invaginations and flaps of membrane observed by FIB-SEM (Figure 2C, Figure 3D). As cells completed downscaling and began to express markers of transition to SPEM (Stage 2 paligenosis; 48-72 hours after injury) and subsequently re-entered the cell cycle, they no longer manifested these contorted apical membranes, although the secreted membranes and glycoproteins (present in the cast-like structures described above) persisted within the lumens of the glands (Figure 1C,D,G,H, Figure S4). The secreted luminal substance was also visible at 24 hours via FIB-SEM (Figure 2B) and in confocal microscopy (Figure S5H) (Brown et al., 2022; Goldenring et al., 2000; Ma et al., 2022; Nozaki et al., 2008). Taken together, we observed an excretory process of material from multiple cellular compartments (at least ER and zymogenic granules) that occurs during cellular downscaling. The material appeared in the lumen via dramatic distortion of the apical membrane into a series of invaginations and interconnecting cavities.

### Excretory process is distinct from autophagy

During cellular downscaling in stage 1 of paligenosis, there is massive upregulation of autophagic and lysosomal structures. (Radyk et al., 2021; Willet et al., 2018). We next investigated whether lysosomes were involved in the excretory process at the apical membrane. We tracked Late Endosomal / Lysosomal LE/Ly) structures by immunostaining for the small GTPase RAB7 and the intrinsic membrane glycoprotein LAMP2 (Figure 5, 6, Figure S7, S8).

At homeostasis, RAB7 marked infrequent, small punctae distributed throughout the cytoplasm (Figure 4A, Figure 5A). After injury, we observed large (up to 5 µm) intracellular RAB7-positive vesicles by 24 hours (Figure 4C, Figure 6A) using optical sectioning. (These are difficult to identify in IHC due to the thickness of a 5-7 µm tissue section and lack of spatial resolution in the enzyme-based IHC assay (e.g., Figure 5B)). We observed that these large, intracellular RAB7^+^ vesicles sequestered a portion of the cellular sulfated glycoproteins (Figure 4D).

**Figure 4.**
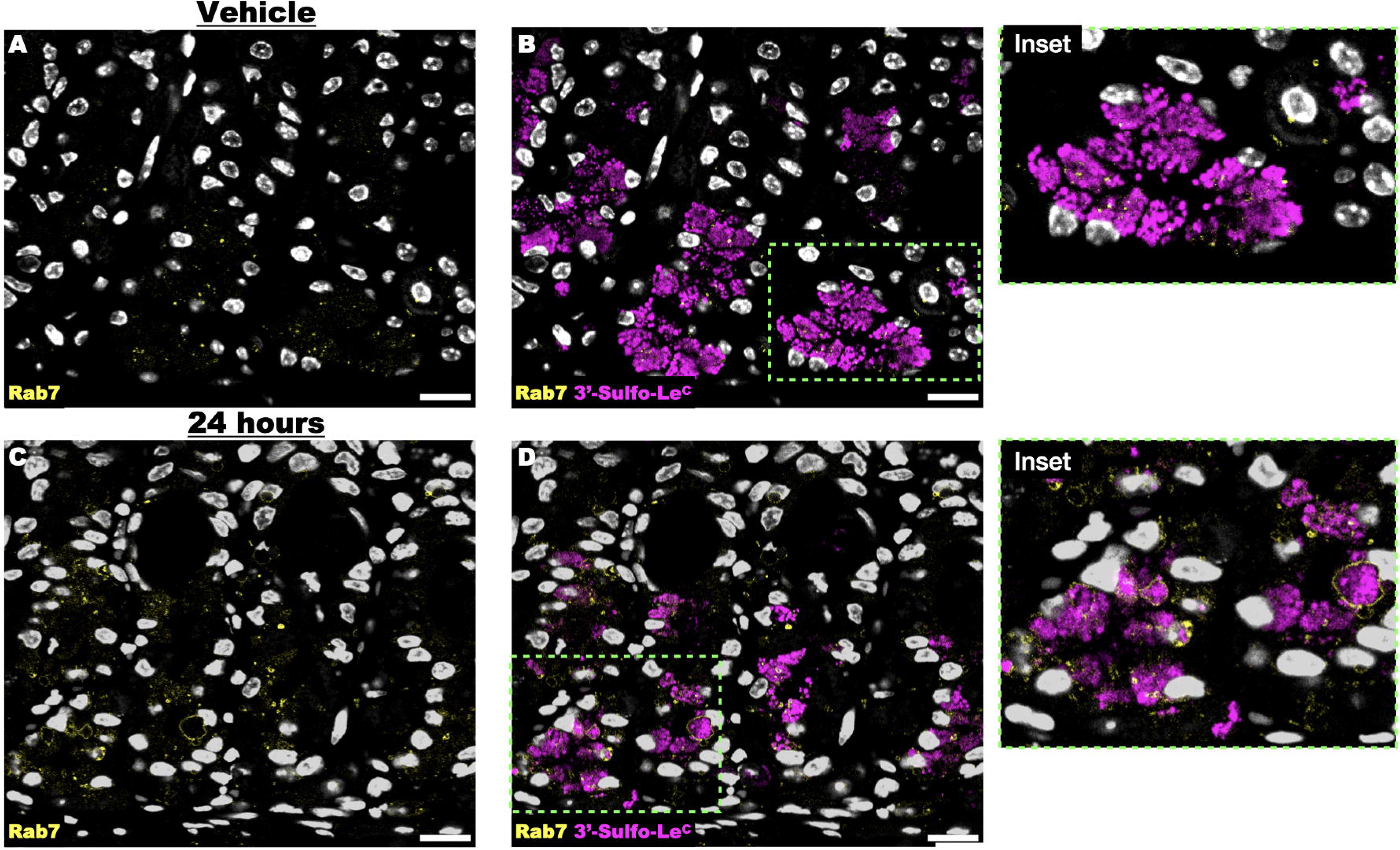
A portion of the sulfated mucins are enveloped by vesicular structures decorated with Rab7. **A,B**. Vehicle-treated stomach demonstrates scattered, small RAB7 positive vesicles, interspersed among zymogenic granules (RAB7 is typically a marker of late endosomes and lysosomes). **C,D**. 24 hours after injury, large RAB7-decorated vesicles envelope sulfated mucins. Pseudocoloring: White: Nucleus (DAPI); Yellow: RAB7; Magenta: 3’-Sulfo-Le^C^ (Das-1). Scale bars = 20 µm.

Large autophagic vesicles can secrete their contents via direct fusion with the apical membrane (Secretory Autophagy, (van Meel et al., 2011)). The defining component of secretion via this pathway is evidence of intrinsic lysosomal markers rerouted to the apical membrane. Despite extensively searching for colocalization of lysosomal markers with the apical membrane during paligenosis we have never observed examples of this pattern in wild-type mice (see, e.g., Figure 6A), suggesting that secretory autophagy is not normally present in chief cells at homeostasis or during paligenosis. The absence of colocalization between lysosomal and apical membrane markers does not absolutely exclude secretory autophagy as the lysosomal proteins could be actively and rapidly reinternalized after secretion.

To determine the role of autophagy in the excretory process occurring in downscaling gastric chief cells, we studied paligenosis in the absence of *Epg5* (ectopic P-granules 5 autophagy tethering factor). EPG5 is a multisubunit tethering complex (MTC) required for fusion of LC3+ vesicles (autophagosomes) with the RAB7+ LE/Ly (Wang et al., 2016). In contrast to wild-type mice, we observed that RAB7 and LAMP2 localized to the apical membrane in the *Epg5^−/−^* mice during paligenotic downscaling (Figure 5B vs D and E vs. F; Figure 6A vs. 6B,C). As would be expected Lamp2 colocalized on the large intracellular vesicles with Rab7 in wild-type mice (Figure S7) and on the apical membrane in *Epg5^−/−^* mice (Figure 6D). Such a localization was observed in paligenosis and not at baseline (Figure 5A-D). In many *Epg5^−/−^*paligenotic cells, the extent of incorporation of LE/Ly membrane into the apical plasma membrane caused the cells to expand their apical membrane, and as a consequence transform the gland lumen into cystic structures (Figure 5F & Figure 6D). These number of cysts in each genotype were quantitated in Figure 5G. EPG5 is a RAB7 effector protein and would be expected to mediate RAB7 localization so, theoretically, loss of EPG5 could lead to RAB7 – which is membrane-associated and not an intrinsic membrane protein – mislocalization to plasma membrane. However, LAMP2 is an intrinsic membrane protein in LE/Ly, so its presence in the apical plasma membrane is most likely due to fusion of LE/Ly with apical membrane, likely with subsequent failure of intracellular retrieval (Figure 6B).

**Figure 5.**
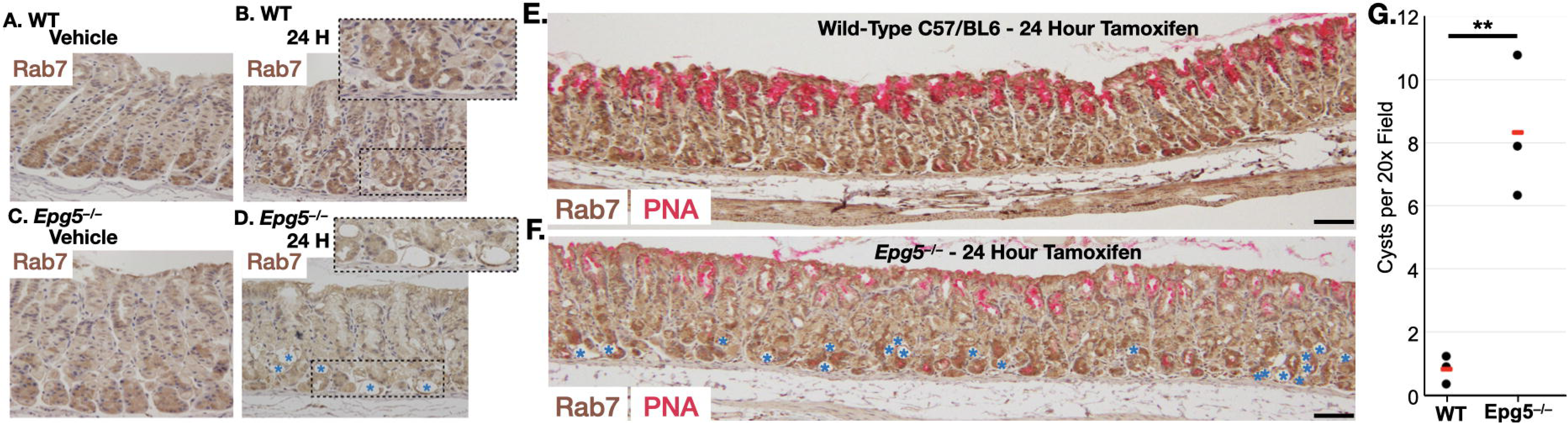
In *Epg5^−/−^* mice, the late endosome / lysosome fuses with the apical membrane during the cellular downscaling reaction. **A,B.** Immunohistochemistry (hematoxylin counterstain) with anti-RAB7 of wild-type C57/Bl6 mouse gastric corpus following (**A**) vehicle treatment or (**B**) 24 hours after injury. **C, D.** Immunohistochemistry of age-matched *Epg5^−/−^* gastric corpus following (**C**) vehicle treatment or (**D**) 24 hours after injury. **A-D**. **E,F**. Wide-field view 24 hours after injury with brown=anti-RAB7 and magenta = PNA (**E**: WT, **F**: *Epg5^−/−^*) demonstrating how tamoxifen injury causes cyst formation in the chief cell zones of *Epg5^−/−^* mice that are mostly absent in wild-type mice. **G.** Quantitation of cystic structures per high power field 24 hours after injury. Black Dots= mean fraction per individual mouse from > 200 fundic glands per mouse, Red Line: Mean of all mice (n=3) per condition depending on experiment. Significance calculated T-Test. Scale bars = 100 µm.

**Figure 6.**
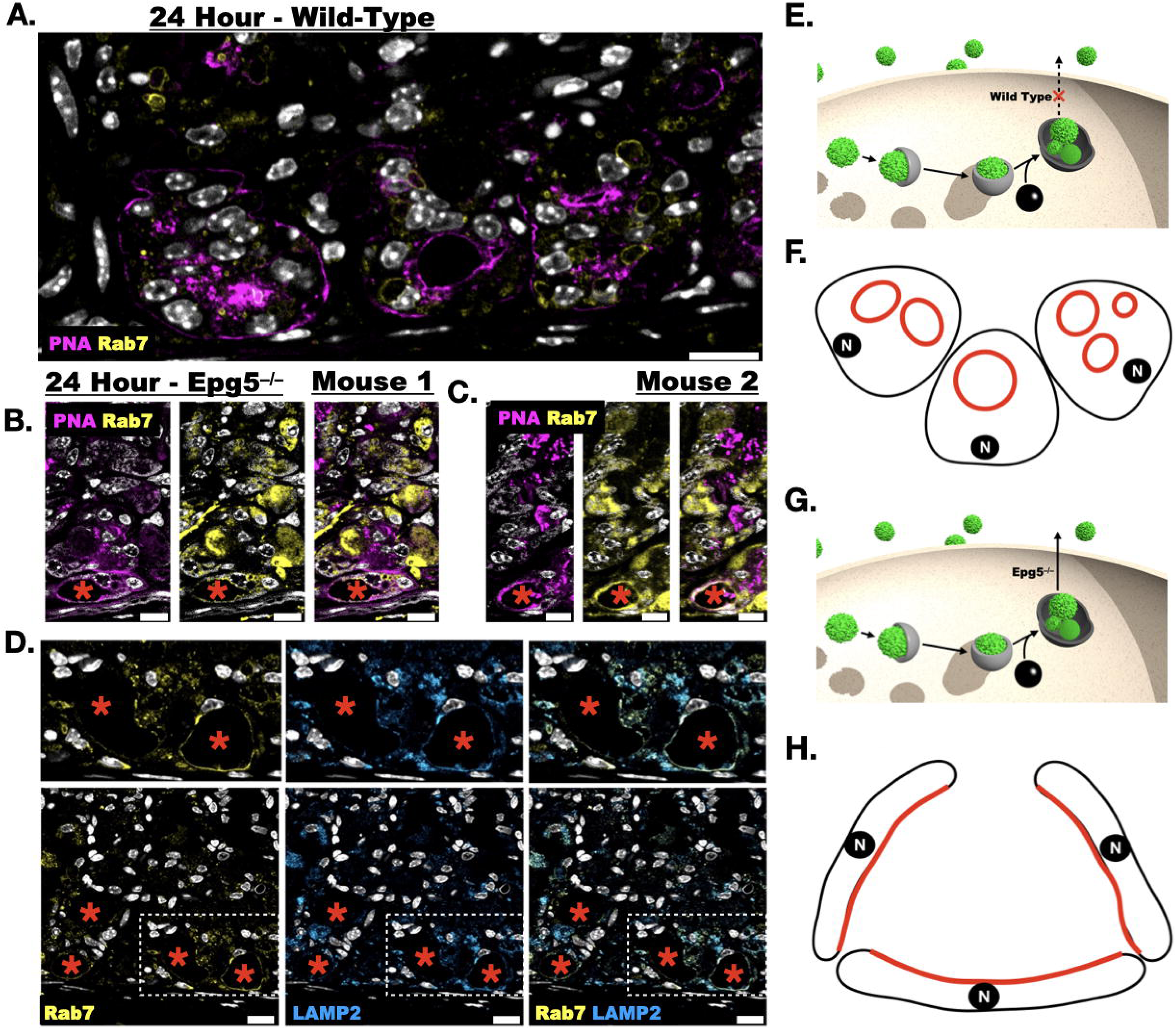
Rab7 does not decorate the apical membrane in wild-type mice but does in *Epg5^−/−^* mice. **A.** At 24 hours after injury RAB7 (yellow) is restricted to vesicular structures but does not decorate the apical membrane (Magenta, PNA lectin). **B, C.** In contrast in the *Epg5^−/−^* background, at 24 hours Rab 7 colocalizes with PNA in the cystic structures. **D.** Further, the intrinsic membrane protein LAMP2 colocalizes with Rab7 in the apices of the cystic *Epg5^−/−^* cells. Same apical distribution with immunohistochemistry in Supplemental Figure 8. Schematic comparing the intracellular trajectory of late endosome / lysosomes in (**E**) wild-type and (**G**) *Epg5^−/−^* mice. (**F,H**) Excessive fusion of lysosomal/autophagic membranes with the plasma membrane without subsequent retrieval causes such expansion of apical membrane that cells expand their luminal surface and flatten into cystic structures. N: nucleus; Black: membrane; Red: late endosome; lysosome membrane. Red asterisks: Cyst like structures. Scale bars = 20 µm.

Summarizing the lysosome and plasma membrane trafficking results: we could detect abundant LAMP2 and RAB7 localization to the apical membrane in *Epg5^−/−^* mice after injury (Figure 5A-F, 6, S8), but we could not detect this localization in wild-type mice, even though we observed numerous distortions of apical plasma membrane associated with the extrusion process at the same timepoints (see Figure 5G for quantified, summed results). Moreover, the cystic structures that arose from the aberrant fusion of the LE/Ly with the apical membrane in the *Epg5^−/−^* background were morphologically distinct from the apical structures in wildtype paligenotic cells, as they did not exhibit the elaborate interconnected chambers that invaginate from the apical membrane in wildtype cells. Thus, we conclude: 1) that secretory autophagy is unlikely to be the principal mechanism responsible for the extrusion of membranous materials we observe in wild-type mice (Figure 6E-H) and 2) that the wildtype, paligenotic apical invagination network is likely a specific cellular structure (e.g., one that would be supported by dynamic cytoskeletal regulation), because the lower energy state is a flat membrane as observed in the *Epg5^−/−^* background.

### Hydroxychloroquine treatment results in intracellular retention of sulfated mucins

We did not observe prominent secretory lysosome activity in wild-type paligenotic cells and next wanted to determine how inhibition of lysosomal function would impact the expression and extrusion of proteins with sulfated mucins. We blocked lysosomal function pharmacologically with hydroxychloroquine, which inhibits vesicular acidification, rendering the lysosomal degradation proteins non-functional, and genetically using mice null for *Gnptab* (N-Acetylglucosamine-1-Phosphate Transferase Subunits Alpha and Beta), encoding a protein essential to trafficking the lysosomal hydrolases to the lysosome.

Treatment with hydroxychloroquine alone reduced the size and apparent number of secretory vesicles at homeostasis (Figure S9). During high-dose tamoxifen-induced paligenotic injury, hydroxychloroquine treatment resulted in retention of sulfomucins on a gland-by-gland basis (Figure 7E). However, since we continue to observe luminal sulfomucins (7B, quantitated in 7F,G), these data suggest that cathartocytosis can still occur in the presence of hydroxychloroquine. The number of injured glands containing secreted antigen was lower with cotreatement of hydroxychloroquine (Figure 7F,G), which may be a consequence hydroxychloroquine limiting the production of this epitope at homeostasis (without injury) (Figure 7E, Figure S9). Consistent with this assertion, we observe the apical deformities characteristic of the excretory cathartocytotic process in hydroxychloroquine treated mice following paligenotic injury (Figure S10). However, there is residual retention of Das-1+ mucins within gastric chief cells (Figure 7B) with tamoxifen + hydroxychloroquine treatment, suggesting incomplete, intracellular clearance of the sulfated mucins and the requirement of ***both*** cathartocytosis and canonical autophagy to downscale cellular machinery *en route* to the cell cycle. In other words, the extrusion of (at least a large portion of the) membranous material does not mechanistically involve lysosomal proteins as expected from absence of apical lysosomal markers in wild-type mice. Additionally, because we never detect lysosomal proteins on the plasma membrane in paligenosis in wild-type C57/Black6 mice, we would expect lysosomes are not directly trafficking most of the extruded membranes into the lumen.

**Figure 7.**
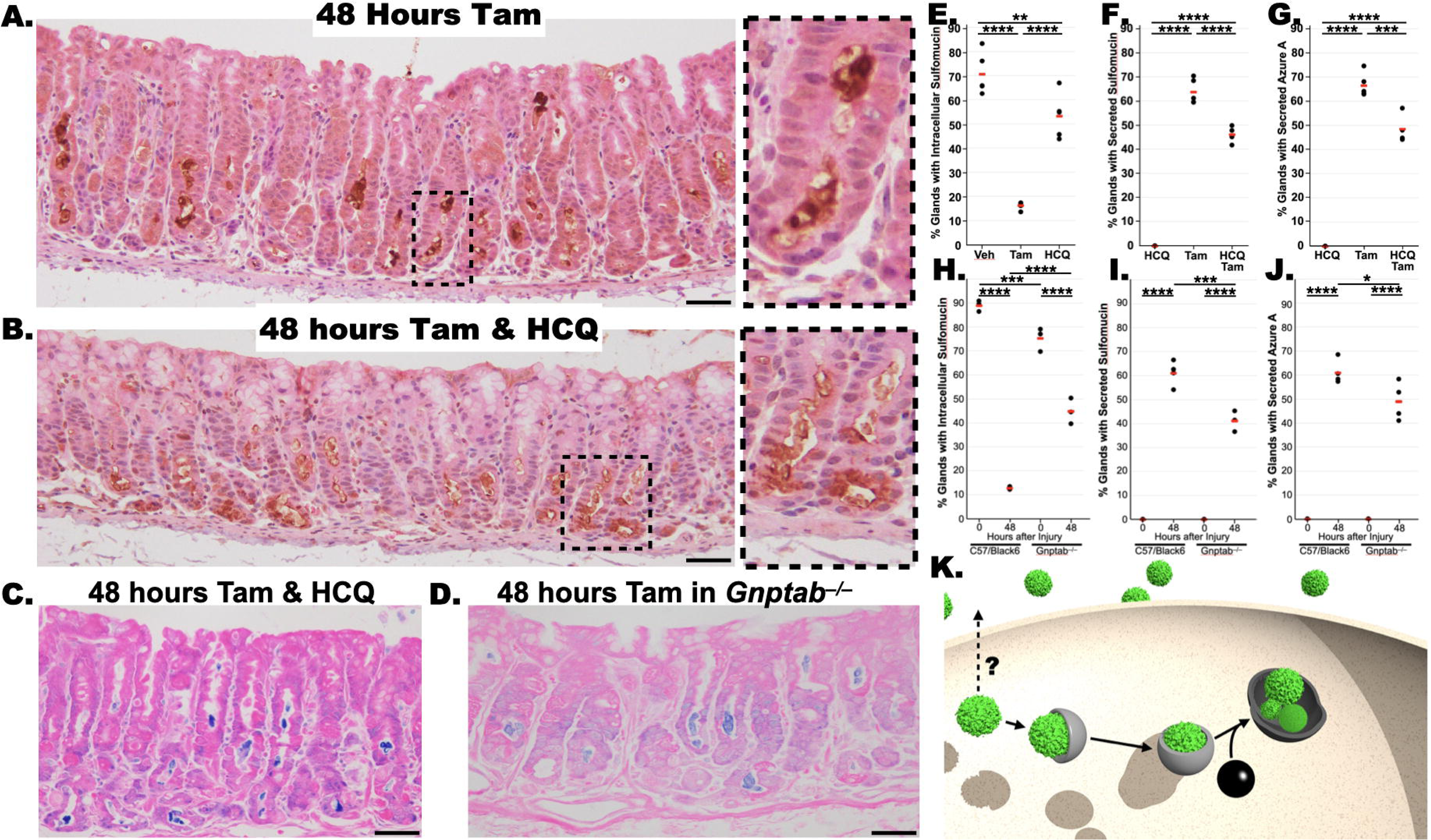
Sulfated mucin excretion requires functioning lysosomes. **A.** 48 hours after high-dose tamoxifen the chief cells are largely devoid of Das-1 positive sulfated mucins (brown; Hematoxylin counterstain, blue) with these mucins being found in the extracellular gland lumen. **B.** Concurrent administration of hydroxychloroquine with tamoxifen results in intracellular retention of sulfated of a portion of the mucins at 48 hours post-injury, a time when sulfated mucins should be absent from the intracellular space. **C.** Azure A (blue; eosin counterstain, pink) reactive material is still secreted after concurrent administration of tamoxifen and hydroxychloroquine. **D.** Secretion of Azure A (blue; eosin counterstain, pink) reactive material in the gland lumen of *Gnptab^−/−^* mice, which do not have functional lysosomes. Scale bars = 50 µm. Wild-type controls for panels C & D are available in Figure 1H. **E**. Quantification of the percent of glands with chief cells harboring intracellular sulfated mucins for hydroxychloroquine alone, tamoxifen, or hydroxychloroquine and tamoxifen. **F,G.** Quantification of gland lumens containing secreted sulfomucins or azure A avid material for hydroxychloroquine alone, tamoxifen, or hydroxychloroquine and tamoxifen. **H.** Quantification of the percent of glands with chief cells harboring intracellular sulfated mucins for wild-type or *Gnptab^−/−^*after vehicle or 48 hours after injury. **I,J.** Quantification of gland lumens containing secreted sulfomucins or azure A avid material for wild-type or *Gnptab^−/−^* after vehicle or 48 hours after injury. Black Dots= mean fraction per individual mouse from > 200 fundic glands per mouse, Red Line: Mean of all mice (n=3-5) per condition depending on experiment. Significance calculated with one-way ANOVA of means with Tukey’s multiple comparison test. Schematic representation depicting secretion is mechanistically independent of canonical autophagy that occurs following injury. Grey: classical autophagy; Black: Lysosome; Green: Generic cytoplasmic organelle (granule, ER, mitochondria, etc.).

### FIB-SEM demonstrates the apical invaginations have phagophore-like activity

In further characterizing the apical membrane deformations occurring during stage 1 of paligenosis, we observed phagophore (Cup)-shaped structures emanating from the apical invaginations (Fig. 8A, Supplemental Movie 3). Additionally, we observed direct fusion of a zymogenic granule with the apical invaginations. At sites of fusion, we observed (1) gradients of electron density suggesting spillage/leakage of the vesicular content and (2) generation of extracellular membranous material. Thus, the excretory process is distinct from simple fusion of the vesicle with the apical membrane (Fig. 8B, Supplemental Movie 4) because the membranous components with electron lucent interiors generated from the fusion event, which are observed throughout the invaginations and in the gland lumen, would not be expected to arise with simple fusion of the vesicle with the apical membrane.

**Figure 8.**
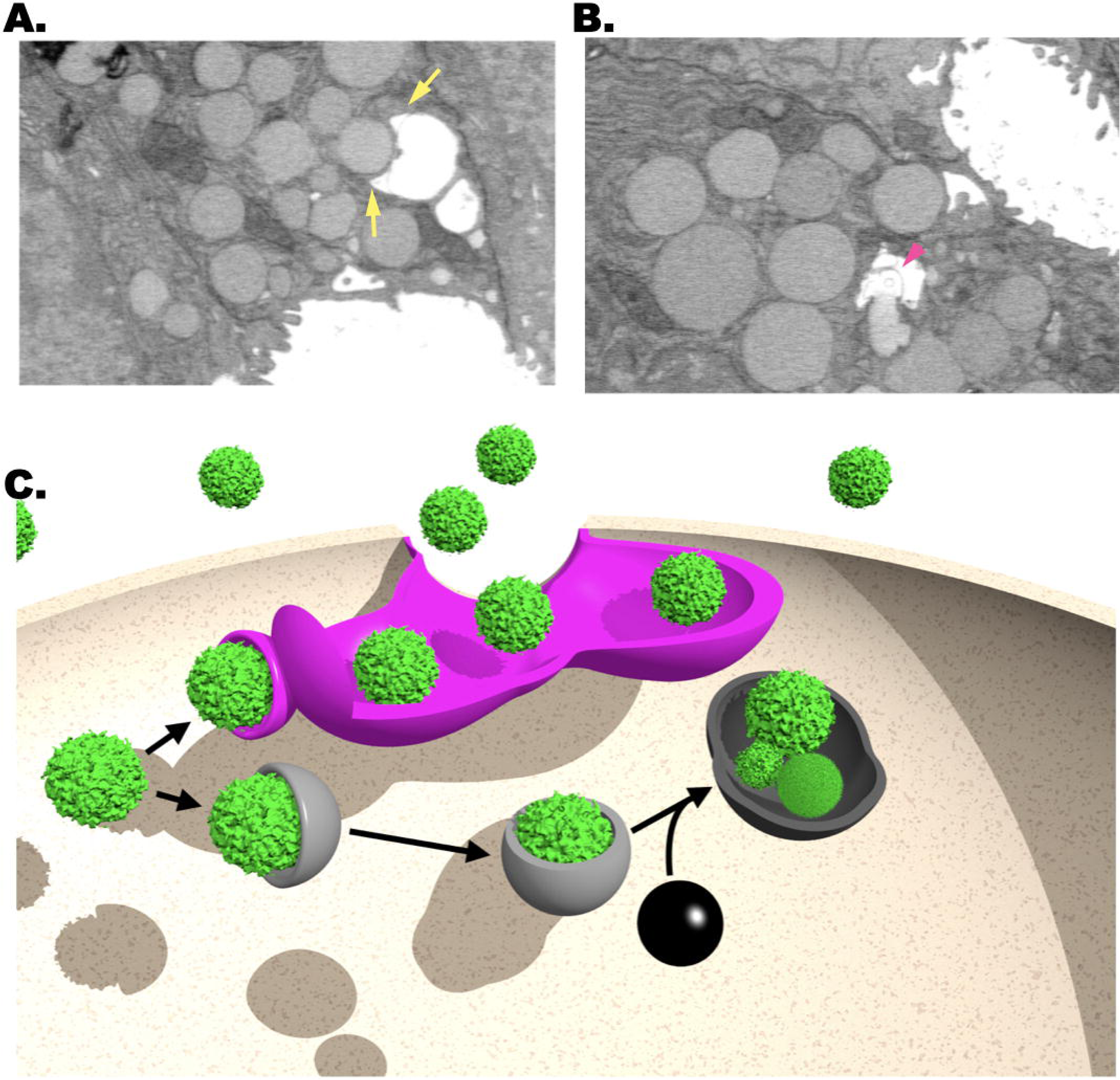
Cathartocytosis Model. **A.** Single 2-dimensional sections from FIB-SEM of chief cell apical plasma membrane at 24 hours post injury demonstrates phagophore-shaped structures (i.e., similar to emerging, double-membrane autophagosomes) contiguous with the apical invaginations (Yellow Arrows). Supplemental Movie 3 shows this section in 3-dimensional context. **B.** Fusion and of a zymogenic granule into an apical invagination with release of cargo as well as membrane into the lumen (Pink Arrowhead). Supplemental Movie 4 demonstrates the section in 3-dimensional context. **C**. Schematic representation of canonical autophagy versus our proposed cathartocytosis process. The two processes occur concurrently after injury and both likely combine to play a role in cellular downscaling. Magenta: Cathartocytosis; Grey: classical autophagy; Black: Lysosome; Green: Generic cytoplasmic organelle (granule, ER, mitochondria, etc.).

## DISCUSSION

Using a synchronous *in vivo* murine model of gastric injury-induced metaplasia, we describe how mature, differentiated cells downscale their cellular machinery *en route* to a proliferative phenotype. Our work supports the importance of lysosomes and autophagic machinery in cellular downscaling (Figure 7), as has previously been reported by our group (Brown et al., 2022; Radyk et al., 2021; Willet et al., 2018). However, we also uncover a cellular process that occurs concurrently with, but does not utilize, autophagic machinery. We propose the term cathartocytosis [Greek: Cellular Cleansing] for this cellular process that bestows a previously undescribed excretory capacity of cellular structures through the apical membrane.

Cellular plasticity is paramount for resurrection of the cell census and maintenance of barrier function following substantial tissue injury (Horowitz et al., 2023; Zuo et al., 2020). Cellular reprogramming to a regenerative state can supersede the homeostatic functions of differentiated cells when injury is severe enough to potentially compromise the barrier. This is especially true for the gastrointestinal epithelium which plays an exceedingly important barrier function protecting the human body against ingested and endogenous caustic, toxic, and pathogenic material to which it is constantly exposed.

Our group has previously described a stereotyped series of cellular-molecular events that occur in differentiated cells following injury to license these cells to proliferate. This process, called paligenosis (Willet et al., 2018), results in a differentiated cells downscaling their cellular machinery to reenter the cell cycle. Here, we studied how these dramatic phenotypic/architectural come about.

An essential early step in paligenosis is the increase in autodegradative machinery, upregulation of the transcription factor *Atf3*, which, in turn, induces expression of *Rab7* (Radyk et al., 2021). Here, we demonstrate that canonical autophagy is responsible for removal of some but not all of the intracellular mature differentiated features and that the cell’s autodegradative machinery is supported by the direct excretion of cellular organelles through the apical membrane to help achieve rapid cellular downscaling (Figure 8C). Further, our analysis indicates that such extrusion is an active secretory process as opposed to simply sloughing of glandular material membranous material as was previously proposed (Goldenring et al., 2000).

The cytoplasmic compartment of the gastric chief cells is mostly occupied by an expansive rough endoplasmic reticulum (ER) and zymogenic granules localized apical to the nucleus (Figure 2A, Figure S5). While, as we previously have reported, some of the ER and granules are indeed degraded by lysosome-dependent machinery, we were also able to show that much of these organelles were jettisoned into the apical lumen by a process we term cathartocytosis. We do not believe what we observe is exosome secretion, because exosomes are characterized as a relatively uniform population of vesicles secreted by fusion of the multivesicular body with the apical membrane (Jeppesen et al., 2023). Further, by transmission electron microscopy, exosomes are approximately as electron-dense as the cytosol because they are formed intracellularly and secreted following fusion of the multivesicular body with apical membrane. In contrast, the excreted vesicular matter we observe is electron-lucent. We can observe a gradient of decreasing electron density as the secreted substance nears the gland lumen (Figure 8B, Movie S4). Lastly, exosomes are formed from invagination and sequestration of cytoplasmic material into the multivesicular body, which then fuses with the apical membrane. As the glycosylation epitopes on mucins are never exposed to the cytosol, the only way for mucins to end up as cargo in the much smaller exosomes would be for the very large (1 μM) secretory mucin vesicles to be enveloped by the multivesicular body, which would then fuse with the apical membrane; however, this would result in the secretion of double membranes (exosome and secretory vesicle inside), which we do not observe (Figure 8). We speculate that the extracellular membranous material we observe being extruded is variegated and vesicle-like because the membranes: (1) were partially lysed during their generation, (2) have fenestrations, (3) formed secondarily in the extracellular milieu from jettisoned lipid material, and/or (4) formed around extraluminal material. Based on the ultrastructural characterization here, we favor one of the first two mechanisms (Figure 8B, Movie S4).

The secreted material appears to be released following docking with phagophore-shaped structures arising from the apical membrane of vesicles independent of a multivesicular body (Figure 8A). The secreted vesicular material we observe is distinct from oncosomes, microvesicles, ectosomes, apoptotic bodies, and exophers as the membrane topology of all these vesicular structures involve budding from the apical membrane (Jeppesen et al., 2023). Other secretory mechanisms have been described involving fusion of organelles containing other vesicles like secretory lysosomes (Maricchiolo et al., 2022; van Meel et al., 2011); however, we have ruled out a significant contribution from secretory lysosomes in wild-type mice using the *Epg5* null mice as positive controls to show we have the tools to detect this process, yet do not observe it in wild-type paligenotic cells. Further, we are not aware of another secretory mechanism that involves the development of an elaborate invagination network of the apical membrane used to release the cellular contents. Lastly, we observe cathartocytosis to occur as a means of rapidly downscaling cellular contents and size, while other secretory processes have been described while roughly maintaining of cell size and contents.

During times of profound exocytosis as we have described here and which occurs in other cell types like those undergoing apocrine secretion, limiting the incorporation of excessive membrane to the apices of the cell is of paramount importance to maintaining homeostatic cell shape and size. For example, in *Drosophila* during exocrine secretion from the larval salivary gland it was found that following vesicle fusion with the apical membrane the remanent membrane is not incorporated into the apical membrane, but instead is supported by actomyosin cytoskeleton and subsequently crumples and removed by endocytosis (Kamalesh et al., 2021). Here, we describe an elaborate set of invaginations from the apical membrane that is likely also supported by the cytoskeleton. The elaborate apical invaginations described here are distinct from vesicle crumpling in *Drosophila* because the size is much larger than a single secretory vesicle, and vesicle crumpling was shown to be a means of eliminating excess membrane from the apices by endocytosis, while the membrane invaginations in cathartocytosis serve as an excretory apparatus.

At a cellular level, the function of EPG-5 was originally identified in a *C. elegans* autophagy screen where it was suggested that it played a role during late stages of autophagocytic degradation (Tian et al., 2010). Consistent with this role, it was later shown in cell lines that *EPG5* functions as a RAB7 effector, augmenting SNARE-mediated fusion of the late endosome with the LC3+ autophagophore (Wang et al., 2016). In the absence of EPG5, these authors found that RAB7 vesicles were promiscuous and fused with other compartments including early endosomes. Here, we found that following injury in the absence of EPG5, the RAB7+/LAMP2+ LE/Ly fused with the apical membrane, an outcome we have not seen before in paligenosis in any other context (Figure 5, 6, Figure S8). Moreover, the *Epg5* null paligenotic cells demonstrate far more dramatic reduction in apical – basal distance / thinning out of the cells, likely a consequence of (1) addition of LE/Ly membranes to the apical domain as well as (2) cell death resulting in stretching of remaining viable cells. Fusion of the LE/Ly with the apical membrane occurs at homeostasis in the osteoclast and is important for bone resorption and described as secretory lysosomes (van Meel et al., 2011). Thus, it appears that EPG5 plays an important role in limiting excessive incorporation of membrane into the apices of the cell presumbably by limiting inappropriate SNARE complexes from forming. It remains to be determined whether such aberrant trafficking of the LE/Ly in *Epg5* null cells is responsible for the known immunodeficiency in humans with Vici syndrome, which is characterized by mutations in *EPG5*. The dramatic phenotypic difference between injured wild-type cells which are cuboidal with stable apical invaginations and injured EPG5^−/−^ cells which are pancake-shaped demonstrate the importance of (1) regulating what membrane compartments are incorporated into the extracellular membrane (2) cytoskeletal support of secretory structures / invaginations.

The fact that sulfated glycoproteins (likely mucins, Figure S2) dynamically redistribute during metaplasia may explain why sulfated mucin expression is heterogeneous / variegated when viewed in chronic metaplasia in humans (Das and Brown, 2023). Such metaplasia would be subject to chronic inflammation and repair in a non-synchronous fashion, so that we might observe cells at various states of sulfated mucin trafficking, as has been reported by us and others in: Barrett’s esophagus (Brown et al., 2021), Intestinal Metaplasia of the stomach (Bodger et al., 2003; Brown et al., 2021), as well as pancreatic intraepithelial neoplasia 3 (Das and Brown, 2023; Das et al., 2021).

We show here that murine zymogenic granules contain 3’-Sulfo-Le^A/C^ mucins (Figure 1) at homeostasis. They also harbor damage-associated molecular patterns (DAMPs) like cathelidicin (LL-37/hCAP8). Both sulfated mucins and cathelidicin bind the pathogenic organism that causes chronic metaplasia and increases risk for gastric cancer in hundreds of millions of people worldwide: *H. pylori* (Hase et al., 2003; Nuding et al., 2013; Veerman et al., 1997a). The binding has been best worked out for sulfated mucins, where they associate with neutrophil activating protein (HP-NapA) in a pH-dependent fashion (Namavar et al., 1998; Teneberg et al., 1997; Veerman et al., 1997a). The importance of HP-NapA as a virulence factor is illustrated by the protective effect of vaccination with HP-NapA prior to *H. pylori* challenge (Satin et al., 2000). Thus, cathartocytosis may serve not just as a means of helping cells downscale rapidly in paligenosis but as a way to help bind and flush *H. pylori* out of the gland lumen. Other defensins are over expressed and secreted in response to *H. pylori* and may also serve similar functions (Boughan et al., 2006; Otte et al., 2009; Patel et al., 2013; Sigal et al., 2019).

In addition to binding *H. pylori*, the 3’-Sulfo-Le^A/C^ laden mucins also bind several host immune receptors (Das and Brown, 2023). For example, 3’-Sulfo-Le^A^ is the most potent ligand for E- and L-selectins (Galustian et al., 1997; Galustian et al., 1999; Green et al., 1992; Wang et al., 2022; Yuen et al., 1994; Yuen et al., 1992). The cysteine-rich domain of the macrophage mannose receptor as well as the dendritic cell immunoreceptor also have affinity towards 3’-Sulfo-Le^A/C^ (Bloem et al., 2014; Leteux et al., 2000). Thus, the secretion of sulfated mucins following injury that causes metaplasia or occurs in the cancer microenvironment could alter immune response.

Note that there are no model systems to study paligenosis or the cellular steps in *transition* from normal, post-mitotic epithelial cells into a proliferating, metaplastic state. If cathartocytosis is unique to paligenosis (and maybe even unique to paligenosis in certain contexts like when pathogens help trigger it), this may be why it has not been observed in cells in culture. Indeed, the highly elaborate basal-apical trafficking change with highly structured, multi-chambered apical plasma membrane might depend on essential contributions from adjacent epithelial cells, mesenchyme (Lee et al., 2023), basement membrane (Bertaux-Skeirik et al., 2017; Khurana et al., 2013), and the immune system (Bockerstett et al., 2020; De Salvo et al., 2021; Jeong et al., 2021; Meyer et al., 2020; Petersen et al., 2018; Petersen et al., 2014). Unfortunately, we are currently limited in our study of the dynamic trafficking changes in cathartocytosis in living cells because we must work in fixed tissue. Developing such *in vitro* model of paligenosis would allow us to approach cathartocytosis (and other processes) using high-resolution and time lapse microscopy and easily overexpress or knockdown key components to determine the underlying mechanics of this cellular process.

Overall, here we describe the membrane shuttling trajectories used by injured cells to downscale their cellular machinery *en route* to metaplastic phenotype. Our data reinforce that canonical autophagy serves an important route, but also uncovers a previously undescribed cellular process that we call cathartocytosis and which bestows excretory capacity to the apical membrane that may help cells not only downscale to a smaller, progenitor-like state during paligenosis and regeneration but also may serve as an anti-microbial mechanism or immunomodulatory function. In the future, we hope to explore if cathartocytosis is common to paligenosis (which is broadly conserved across organs and species) and/or if it also occurs in injured cells that are not undergoing massive activation of cell plasticity.

## MATERIALS & METHODS

### Animal studies and reagents

All experiments using animals followed protocols approved by the Washington University in St. Louis, School of Medicine Institutional Animal Care and Use Committee. WT C57BL/6 mice were purchased from Jackson Laboratories (Bar Harbor, ME). *Epg5^−/−^* mice were obtained from breeding *Epg5^+/−^* mice, a kind gift from Megan Baldridge (Lee et al., 2022; Zhao et al., 2013a; Zhao et al., 2013b). Likewise, *Gnptab*^−/−^ were obtained by breeding *Gnptab*^+/−^ mice (A kind gift from Stuart Kornfeld).

Tamoxifen powder (Toronto Research Chemicals) was initially solubilized in 100% ethanol via sonification after which it was emulsified in sunflower oil (Sigma-Aldrich) at a 9 Oil :1 EtOH ratio(Saenz et al., 2016). Tamoxifen (5 mg/20 g body weight; Toronto Research Chemicals) was injected intraperitoneally daily for up to 2 days or until mouse was euthanized for histologic examination. Hydroxychloroquine (120 mg/Kg by intraperitoneal injection) was administered 24 hours prior to the first tamoxifen injection and then with all subsequent tamoxifen injections until the tissue was harvested. All mouse experiments were performed on mice aged 6-10 weeks.

### Imaging and tissue analysis

Following anesthetizing with isoflurane and cervical dislocation, murine stomachs were excised, flushed with phosphate buffered saline and fixed overnight with 10% formalin. They were washed and equilibrated in 70% ethanol for several hours prior to embedding in 3% agar and routine paraffin processing. Sections (5-7 μM) were prepared for immunohistochemistry and/or immunofluorescence by deparaffinization using Histoclear and an alcohol series for rehydration. After which antigen retrieval was performed in 10 mM citrate buffer, pH 6.0 in a pressure cooker. Tissue was blocked with 2% BSA and 0.05% Triton X-100. Primary and secondary antibodies were diluted in 2% BSA and 0.05% Triton X-100. Vector ABC Elite kit was used for immunohistochemistry. Immunohistochemistry slides were mounted with Permount and immunofluorescence with Prolong Gold. Brightfield images were taken on either a Nanozoomer (Hamamatsu 2.0-HT System) for quantitation or Olympus BX43 light microscope. Confocal images were obtained on a Zeiss LSM880 confocal microscope.

### Azure A staining

Slides of paraffin embedded tissue were dewaxed and rehydrated with an ethanol series to 30%. Antigen retrieval must not be performed as this results in loss of staining. Slides were then stained with 0.1% Azure A Chloride in 30% ethanol for 1 minute. Slides were then washed in water. Placing slides in citric acid / sodium phosphate buffer, pH 4 results in royal blue nuclear staining and dark blue cytoplasmic staining. The slides were then transferred to 95% ethanol and stained with Eosin Y. The slides were subsequently mounted with permount.

### Focused-ion beam scanning electron microscopy

The tissue fixation and acquisition parameters have been previously described (Lo et al., 2017; Radyk et al., 2021). Three-dimensional reconstructions of either the gastric chief cells or the gland lumen were obtained by initially masking using ilastik-1.4 using the carve protocol (Berg et al., 2019). These masks were exported to FIJI (ImageJ) (Schindelin et al., 2012) to crop and create an image stack. This image stack was then imported to ChimeraX (Pettersen et al., 2004) to generate and render the reconstruction. Movies form the rendered images with iMovie.

### Western blot

Western blot samples were prepared using ∼30 μg of protein in standard SDS-PAGE lammeli buffer with 5% β-mercaptoethanol. Samples were heated to 70 °C for 10 minutes prior to running through NuPAGE precast gels. Protein was transferred in NuPAGE transfer buffer containing 20% methanol to nitrocellulose which was blocked using 5% BSA in PBS. Nitrocellulose membrane was incubated with primary antibodies in blocking buffer overnight at 4 °C. Membranes were washed with PBS and then incubated in appropriate secondary antibodies at 1:10,000 dilution in blocking buffer. After extensively washing, membrane was imaged using LiCOR Odyssey.

### Dot blot

Sample was applied to nitrocellulose using a BIORAD vacuum apparatus, after which the nitrocellulose was stained and imaged in an identical fashion to western blots.

### Chloroform extraction

Flowchart is displayed in Supplemental Figure 2D. 24 hours after a C57BL/6 mouse was injected with tamoxifen (See above), the 50 mg of stomach tissues was excise and homogenized in 0.5 mL of phosphate buffered saline w/ protease inhibitors. To 0.2 mL of this homogenate, 1.2 mL of methanol and 2 mL of chloroform was added and incubated at 37C in a shaker. An additional 1 mL of methanol was added, and the mixture was spun at 2000g for 30 minutes at 20C. The majority of the supernatant was removed; however, since the pellet was still soft it was spun at 20,817g for an additional 2 minutes and the remaining supernatant removed. The pellet was resuspended in 2 mL of 1:2:0.8 (chloroform:methanol:water) and 10% of the solution was banked for dot blot analysis (Pellet #1). The remaining solution was shaken at 37C for 2 hours. After which the slurry was spun at 20,817g for 2 minutes. The supernatant was removed and added to the prior supernatant and the pellet banked for analysis (Pellet #2). The supernatant and both pellets were dried prior to be resuspended in 4% SDS (supernatant and Pellet #1) or 4% SDS and 6M Urea (Pellet #2). Equivalent volume percentages were loaded for dot blot analysis.

### Deglycosylation

Mucins in solution were treated with O-Glycosidase, Neuraminidase, or PNGaseF according to manufacturer’s / New England Biolab’s instructions.

Slides of paraffin embedded tissue were dewaxed and rehydrated, followed by antigen retrieval as described above. To remove O-linked glycans the sections were incubated in 0.1N NaOH for 2 hours at 20C (Longer incubations or higher temperatures result in tissue being dislodged). The slides were then transferred to phosphate buffered saline and assayed as described above. N-linked glycans were removed by treating with PNGaseF for 2 hours at 37C. Slides were also treated with O-glycosidase and neuraminidase for 2 hours at 37C.

### Methods

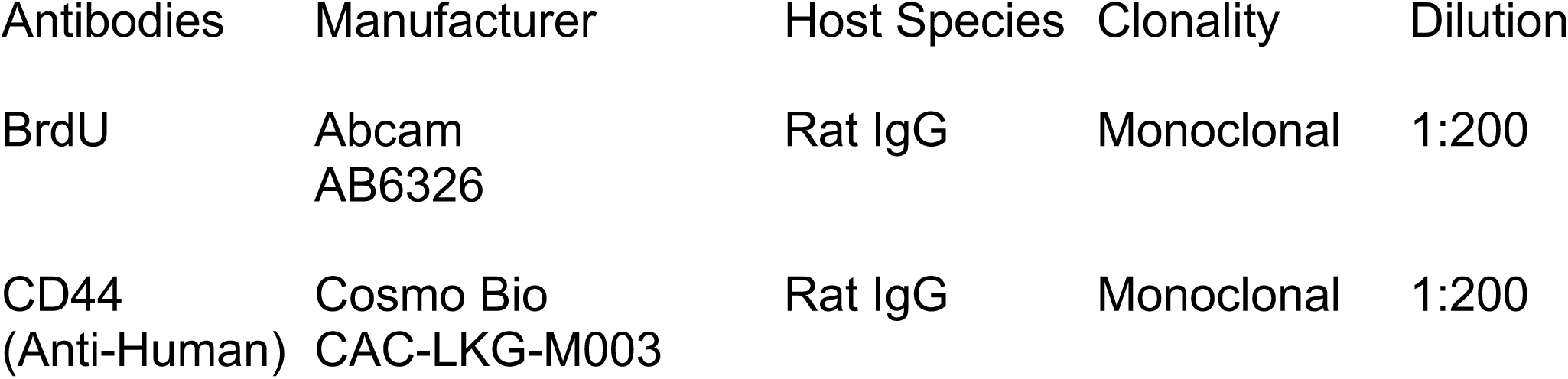
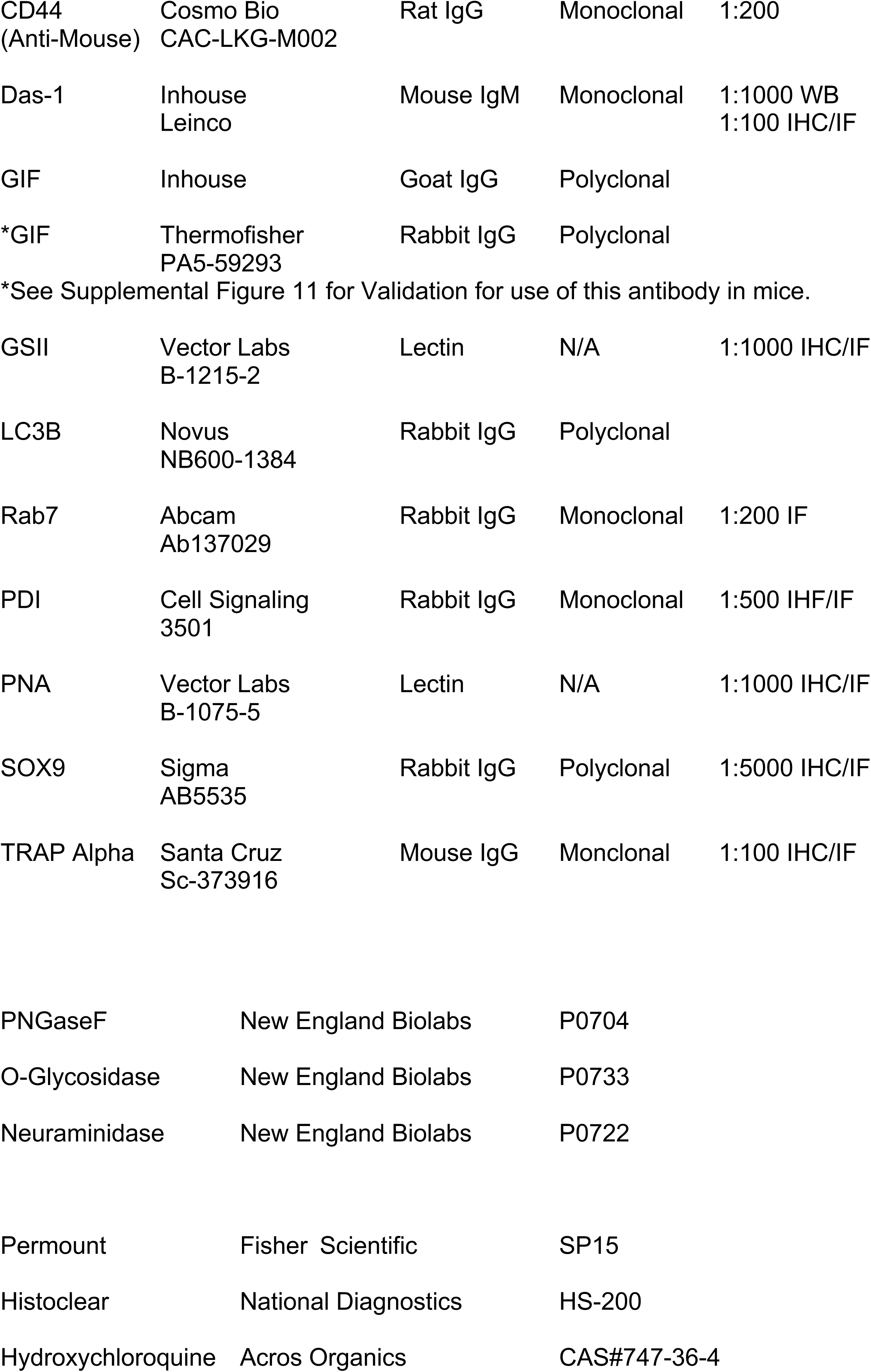
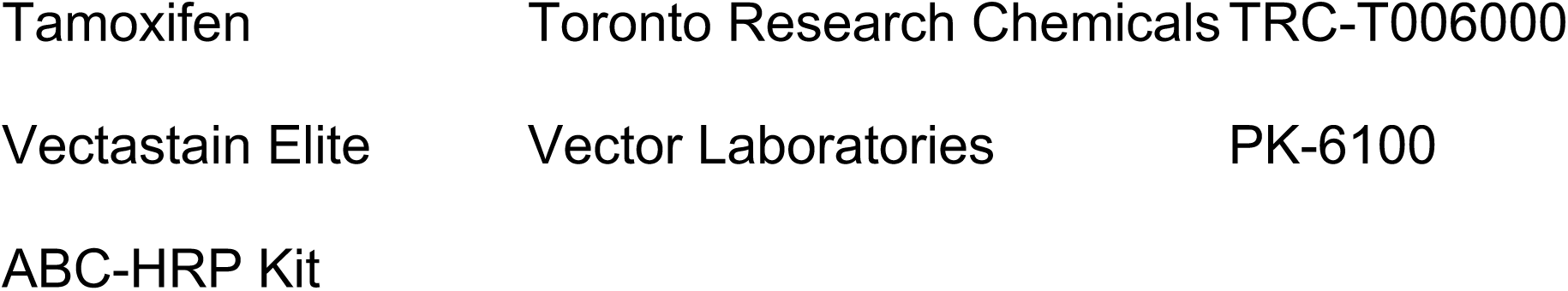

## ACKNOWLEDGEMENTS

Jeffrey W. Brown is supported by NIH K08 DK132496, Department of Defense, through the PRCRP program under Award No. W81XWH-20-1-0630, the American Gastroenterological Association AGA2021-5101, R21 AI156236, P30 DK052574. We thank Stuart Kornfeld for providing the *Gnptab^−/−^* allele and Megan Baldridge for providing the *Epg5^−/−^* allele.

## Author Contributions

Study Concept and Design: JWB, XL, JCM. Data Acquisition and Analysis: JWB, XL, XL, GN, TN, MDR, JB; Drafting the manuscript: JWB, MDR, JCM. Revisions to the manuscript: JWB, XL, GN, TN, MDR, JB, JCM.

---

**Supplemental Figure 1.** Schematic representation of relevant Lewis glycans. Type 1 differs from Type 2 in the arrangement of the galactose and fucose about the N-Acetylglucose. The antibody Das-1 recognizes 3’-Sulfo-Le^A^ and 3-Sulfo-Le^C^, the difference between the glycans being 3’-Sulfo-Le^C^ lacks a fucose. 3’-Sialyl-Le^A^ is the oncoantigen CA19-9.

**Supplemental Figure 2.** 3’-Sulfo-Le^A/C^ is present on large O-linked glycoproteins. Concentrated conditioned media from containing LS174T cells has been used as the standard for 3’-Sulfo-Le^A/C^ substance (Brown et al., 2021). **A**. Western blot of 3’-Sulfo-Le^A/C^ standard showing NaOH treatment results in loss of most signal. **B**. Western blot of 3’-Sulfo-Le^A/C^ standard following treatment of PNGase F and O-Glycosidase with and without Neuraminidase. No change in Das-1 intensity was noted in any of the conditions; however, note that PNGase likely shifts Das-1-labeled bands to lower molecular mass due to loss of N-linked glycans elsewhere on the protein backbone (arrowhead). Note also that the recombinant PNGaseF enzyme added to these reactions also labels with Das-1 (arrow and see next panel).. **C.** At the high concentrations of recombinant enzymes recommended by the manufacture to use in these deglycosylation assays, we observed Das-1 reactivity on the recombinant enzymes run in lanes by themselves. Note both PNGaseF and O-Glycosidase (pure reagent as supplied by NEB) bind Das-1 unless they are treated with NaOH. It is unclear whether this is due to these recombinant proteins expressed in bacteria containing 3.-Sulfo-Le^A/C^ epitopes or non-specific binding with proteins at high concentrations. In sum, the removal of the glycan with sodium hydroxide, but not O-glycosidase is consistent with 3’-Sulfo-Le^A/C^ residing on a branched O-Linked mucin. **D.** Protocol flowchart for chloroform extraction. **E**. Dot blot of PBS alone, first supernatant, first pellet, second supernatant, second pellet. The signal in the aqueous (pelleted) but not chloroform (supernatant) fractions demonstrates 3’-Sulfo-Le^A/C^ is on a glycoprotein and not hydrophobic moiety like a lipid. **F, G**. Immunohistological confirmation that Das-1 reactivity is lost following (**G**) treatment with NaOH relative to control (**F**). Scale bars = 50 µm.

**Supplemental Figure 3.** Representative fluorescence micrograph demonstrating proximity of gastric intrinsic factor (GIF) and 3’-Sulfo-Le^A/C^ (Das-1 antibody) 8 hours after injury. The 8 hour time point was chosen in the unlikely event that injury induced a new population of secretory vesicles in paligenotic cells. The similar subcellular location of GIF (purple) and the Das-1 epitope (cyan) demonstrate that sulfated mucins are associated with chief cell secretory vesicles at all times in the metaplasia cascade. Scale bars = 20 µm.

**Supplemental Figure 4.** Azure A stains the endoplasmic reticulum of the gastric chief cell and permitted tracking of this compartment during paligenosis. Azure A histological stain highlights subcellular changes occurring within chief cells undergoing paligenosis. (**A,E**) Vehicle-treated wild-type mice demonstrated strong dark blue basal staining tapering towards cell apex where zymogenic granules that were not Azure A avid localize. Chief cell staining paralleled confocal microscopy using anti-protein disulfide isomerase, an established endoplasmic reticulum marker PDI (cf. Supplemental Figure 5). Nuclei were royal (light?) blue due to affinity of Azure A for deoxyribonucleic acids (Supplemental Figure 7). (**B,F**) At 8 hours after injury the intense dark blue basal staining began to dissipate. (**C,G**) At 24 hours after injury, Azure A reactivity filled gland lumens, and the apical-basal distance became smaller. (**D,H**) At 72 hours after injury, the chief cells were small, cuboidal, and had scant cytoplasm. Gland lumens were full of Azure A-avid material. **I**. The synchronicity of the tamoxifen injury model can be appreciated at 72 hours in a lower-magnification image demonstrating all glands full of Azure A-reactive material. As this material labels with ER marker PDI (cf. Supplemental Figure 5), and Azure A stains ER in homeostasis, the lumen material may largely be extruded ER. Given (**A-C**). Scale bars in A-D = 50 µm; E-H = 20 µm.

**Supplemental Figure 5.** Excretion of endoplasmic reticulum markers. IHC during paligenosis (**A**. vehicle, **B**. 8 hr, **C**. 24 hr, **D**. 72 hr) demonstrates secretion of PDI. **E, F**. The Gastric Chief Cell Cytoplasm was primarily composed of Endoplasmic Reticulum (PDI, Yellow) and Zymogenic Granules (Das-1, Magenta). Epitopes marked by these two antibodies are mutually exclusive. The lectin Griffonia simplifolia (GSII) was used as a neck cell marker (cyan). G-H. Epitopes from the ER as well as the secretory granules are excreted following injury. G. Vehicle treated. H. 24 hours after injury. Scale bars in A-D = 50 µm, Scale bars in E-H = 20 µm.

**Supplemental Figure 6.** Demonstration that Azure A binds nucleic acids. Agarose Gel used for genotyping mice stained with Azure A.

**Supplemental Figure 7.** RAB7 (yellow) and LAMP2 (cyan) colocalized on large intracellular vesicles 24 hours after injury in wild-type C57/Black6 mice. Nuclei (via DAPI) are white. Scale bars = 20 µm.

**Supplemental Figure 8.** Distribution of RAB7 in *Epg5* null mice during paligenosis. **A.** In *Epg5*^−/−^ mice, RAB7 was absent from the apical membrane and formed small punctae throughout the cell. This is similar to what was observed in wild-type C57/Bl6 mice (cf Fig. 5A). **B.** RAB7 associated with the apical membrane at 24 hours after injury in the *Epg5* null background, but not in wild-type mice (**D**). **C**. At 48 hours a few glands are still composed of very flat chief cells and larger Rab7 structures are appreciated at this time point compared to (**D.**) Wild-Type stomachs 48 hours after injury. Scale bars = 50 µm.

**Supplemental Figure 9.** Hydroxychloroquine treatment results in smaller zymogenic granules. **A.** Wild-type, vehicle treated mice. **B.** High-power view wild-type, vehicle treated mouse stomach. **C.** Hydroxychloroquine treated mouse stomach. **D.** High-power view hydroxychloroquine treated mouse stomach. IHC performed with the Das-1 antibody, which is reactive against 3’-Sulfo-Le^C^ in murine tissue. **Scale** Bars = 50 µm.

**Supplemental Figure 10.** The dramatic apical deformities are present at 24 hours following injury in hydroxychloroquine treated animals. **A-D.** 4, representative confocal micrographs of tamoxifen-injured hydroxychloroquine treated mice demonstrates identical invaginations (green arrowheads) and membrane flaps (green arrows). Similar to wild-type vehicle? mice and unlike *Epg5^−/−^* there is very little overlap of lysosomal markers (LAMP2) and membrane markers (PNA) in hydroxychloroquine treated mice. Pseudocoloring: White: DAPI/Nuclear; Yellow: Lamp2; Magenta: PNA Lectin. Scale bars = 20 µm.

**Supplemental Figure 11.** Validation that the thermofisher Anti-GIF antibody PA5-59293 is reactive against murine GIF as well. **A-C.** Identical histologic distribution of PA5-59293 (red) and anti-GIF (green, produced by David Alpers). **D-F.** Western blot mouse stomach lysate with vehicle treatment and 24 hours after injury demonstrating comparable bands. The confusion may derive from GIF being present human parietal cells, but murine chief cells. Nonetheless, the antibody PA5-59293 can be used in mouse tissue.

